# Using nano zero valent iron supported on diatomite to remove acid blue dye: synthesis, characterization and toxicology test

**DOI:** 10.1101/755975

**Authors:** Erick Justo-Cabrera, Ernesto Flores-Rojas, Denhi Schnabel, Héctor Poggi-Varaldo, Omar Solorza-Feria, Luz Breton-Deval

## Abstract

The aim of this work was to synthesize and characterize nanoscale zero-valent iron (NZVI) supported on diatomaceous earth (DE) at two different molar concentration 3 M and 4M (nZVI-DE-1 nZVI-DE-2), to test the decolorization treatment of acid blue dye (AB) and perform a toxicological test using zebrafish. The synthesis of the nanoparticles was obtained using the chemical reduction method and the material was characterized by X-ray diffraction, Scanning Electron Microscopy (SEM), Energy-Dispersive X-Ray (EDX), and transmission electron microscopy and Specific Surface Area (BET). The results showed spherical forms in clusters between 20 to 40 nm of zero valent iron supported on diatomaceous earth. The removal of 1 g/l of AB from water treated with NZVI-DE-1 and NZVI-DE-2 reached the decolorization of 90% and 98% of all dye. While controls like NZVI and DE-1 and DE-2 achieved the removal of 40, 37 and 24 % of the dye. Toxicological analysis using zebrafish showed that AB causes a severe defect in development and embryos die after exposure. However, the water samples treated with NZVI-DE-1 and NZVI-DE-2 are not harmful for the zebrafish embryos during the first 24 hours. We conclude that the use of NZVI-DE-1 and NZVI-DE-2 is a promising treatment for dye pollution.

## Introduction

The concentration of dyes in industrial effluents can reach 500 mg/L and pollute local freshwater, reducing the efficiency of sunlight and thereby impeding the process of photosynthesis. (Jung et al., 2016). As a result, the water temperature of the stream decreased, and photoautotrophs organism like algae, euglena, and cyanobacteria could no longer survive (Bide, 2007). The death of these organisms is an ecological loss because they play an essential role in the cycle of nutrients and are capable of absorbing organic matter present in the stream. They can also remove carbon dioxide from the atmosphere, are an indispensable source of food and oxygen for several organisms (Callieri, 2014). Pollution due to synthetic dyes also affects the health of aquatic and terrestrial organisms (Samchetshabam et al., 2017).

Acid blue (AB) was selected for this study because it is used widely by the industrial sector, including for cosmetics, food coloring or for dyeing different fibers, and is commonly mixed with sulfuric acid to make it more soluble before industrial application thereby increasing the toxicity (Ammar et al., 2006). AB is not only toxic for aquatic life but can also cause skin irritation, cornea damage, and promote the development of tumors and cancer (Khelifi et al., 2008).

Several kinds of research have been carried out with the aim of removing dyes from water using aerobic or anaerobic degradation, filtration, adsorption, membrane filtration and other methods (Pacheco-Álvarez et al., 2019; Saleh, 2019; Wang et al., 2018). However, all of these processes have disadvantages such as the extended time required for the treatment, high operational costs, and efficiency (Ziarani et al., 2018).

Recently, nanomaterials have been used to remove several pollutants including dyes such as crystal violet (Răducan et al., 2019), methyl orange (Fan et al., 2009), acid red 88 and Black 5 (Sharma and Shirkot, 2019).

Materials made of nanoparticles have a relatively high surface area to mass ratio rendering them more reactive compared to the conventional materials used for water treatment (Quinn et al., 2005). However, this technology has to overcome some challenges in order to be applied successfully; namely that during synthesis the nanoparticles tend to aggregate easily thereby losing surface area (Raychoudhury et al., 2012; Zhang et al., 2013) and becoming toxic for aquatic life (Krysanov et al., 2010). To avoid these problems, we supported the nanoparticles on diatomite, a well-known filter material in the industry, and completed a toxicological test using zebrafish.

Zebrafish (*Danio rerio*) have been established as an ideal model for toxicological studies to test the effects of contaminants such as alkaloids, glycosides, metals, alcohols, carboxylic acids, among others (Chen et al., 2011; Qiang and Cheng, 2019). Zebrafish have a number of characteristics that make them a good model for testing toxicity. Firstly, female zebrafish are able to produce hundreds of eggs and embryos are transparent which allows for close observation of development under microscope. Secondly, the rapid growth of zebrafish compared to other vertebrates makes it an ideal model for high-throughput analysis (Ali et al., 2011).

Therefore, the objective of this research was to characterize and synthesize nanoscale zero-valent iron supported on diatomaceous earth and to test if treated water had a negative effect on the viability of zebrafish embryos.

## Materials and Methods

### Preparation of diatomite earth and synthesis of nanomaterial

The diatomite earth (DE) was washed with 1 M HCl for 8 h under agitation (150 rpm). After the acid-washed, the DE was rinsed with water, before using as NZVI support. Two types of NZVI-DE were synthesized. The first identified as NZVI-DE-1 was prepared from 0.3 M FeCl_2_.4H_2_O at a 40/60 proportion of FeCl_2_.4H_2_O / DE, while the second type identified as nZVI-DE-2 was made at a 50/50 proportion of FeCl_2_.4H_2_O / DE. NZVI and NZVI-DE-1 and 2 were obtained using the chemical reduction method of FeCl_2_.4H_2_O in aqueous solution using NaBH_4_ as a reducing agent, due to its simplicity and efficiency in securing NZVI (Fu et al., 2014). The FeCl_2_.4H_2_O and DE were added to a previously ethanol deoxygenated by bubbling it with N_2_ gas for 30 min. The iron salt was dissolved in 50 mL ethanol, and the solution was kept under N_2_ bubbling and stirring for 30 min. 400 rpm at 25°C. Later, 1.5 M NaBH_4_ solution was slowly added into the FeCl_2_/DE solution. After the reaction, the solution was kept stirring at 700 rpm for 60 min. In the end, both types of NZVI-DE were washed 10 times with ethanol and dried in an oven, bubbling with argon at 50 °C. The synthesis of the NZVI without DE support followed the same technique.

### Characterization of iron nanoparticles

#### X-Ray Diffraction (XRD)

The XRD analysis was carried out using a Bruker D8 Advance Eco diffractometer coupled with a copper source without a monochromator; the samples were placed in a sample holder with a 2θ range of 5°-130° at a size and time of 0.02° and 0.2 seconds. The analysis of the crystalline phases was carried out using Match software.

#### Scanning Electron Microscopy (SEM) with Energy-Dispersive X-Ray (EDX)

A scanning electron microscope (SEM) HRSEM-AURIGA Zeiss integrated with an X-ray scattered energy (EDX) analyzer at a voltage of 1-10 keV at high vacuum was used. Both the mapping and the elemental compositional analysis were carried out by selecting random areas using the EDX analyzer. The samples were placed on graphite tape supported on metal discs.

#### Transmission Electron Microscopy (TEM)

A transmission electron microscope (TEM) JEM-ARM200F operated at a high vacuum was used to describe the morphology, size, and distribution of the particles. The samples were prepared by dispersing a small amount of powder in absolute ethanol with the aid of an ultrasonic bath. The dispersion (10 μl) was applied on a copper mesh for TEM of 300 mesh with Lacey / Carbon film and allowed to evaporate in a desiccator.

#### Specific Surface Area (BET)

The specific surface area was measured using the Brunauer-Emmett-Teller (BET) method of N_2_ adsorption, using a Micromeritics Gemini 2360 surface area analyzer. Before analysis, the samples were degassed under vacuum at 70 °C for three hours.

#### Batch degradation experiments

The experimental design consisted of a factorial analysis to evaluate two different relationships between the quantity of NZVI and DE (NZVI-DE-1 = 40/60, and NZVI-DE-2 = 50/50) but hold the same quantity of iron (62 mg) in every treatment. The experiment was carried out using serum bottles with 50 mL of water contaminated with dye (1000 mg/L) loaded with 658 mg of Nano NZVI-DE. The controls were (i) DE-1 and DE-2 to evaluate dye degradation to adsorption process (ii) NZVI to assess the effect of free nanoparticles without a support and (iii) AD to calculate the abiotic degradation. The experiments were carried out at 25° C with a mix of 150 rpm. All experiments were performed in triplicate. The removal of AB was measured with the UV spectra (BioSpectrometer, Eppendorf) at 526 λ nm.

#### Fish maintenance and strains

AB line and wild-type zebrafish (*Danio rerio*) embryos were obtained from natural crosses and raised at 28 °C based on standard procedures (Westerfield, 2000). Eggs were obtained by random pairwise mating of zebrafish. The eggs were harvested the following morning and transferred into plastic Petri dishes (60 eggs per dish) containing 10 ml fresh embryo water. Further, unfertilized, unhealthy, and dead embryos were identified under a dissecting microscope and removed. Morphological criteria determined embryonic stages, according to Kimmel and collaborators (Kimmel et al., 1995). Zebrafish were handled in compliance with local animal welfare regulations, and all protocols were approved by the ethics committee (Instituto de Biotecnología, UNAM).

#### Toxicological studies

At 3.5 hours post-fertilization (hpf), embryos were again screened, and any additional dead and unhealthy embryos were removed. Embryos at the sphere stage were selected and transferred into a 48 well plate; ten in each well. Later the embryo water was absorbed, and 300 ul of the treated water samples (AB dye, NZVI-DE-1, NZVI-DE-2, DE-1, DE-2) were loaded into each well; experiments were performed in triplicate. The control was a set of embryos grown with embryonic water in the same plate as well as in a petri dish independently. The embryonic plates were cultivated in a moist chamber at 28 °C from 6 to 24 hpf, and the viability of the embryos was observed under a dissecting microscope. Embryos at 24 hpf were anesthetized with tricaine, immobilized with methylcellulose on agar plates and visualized with a stereomicroscope (Leica MZ 12.5), photographed using a CCD camera (AxioCam MRc 5, Zeiss) and AxioVision Rel. 48 software. Statistical analysis was performed with three independent experimental replicates. The exact number of biological replicates is indicated in the figure legends. Statistical analysis was performed with Prism (GraphPad) Student’s t-test. Error bars in column graphs represent the standard deviation of the mean (s.d.).

## Results and Discussion

### Synthesis and Characterization of iron nanoparticles

The SEM/EDX analysis of DE showed high levels of oxygen at around 52% given that most of the materials that form DE are oxides, such as silicon oxide or aluminum oxide (Table 1).

**Table 1.**
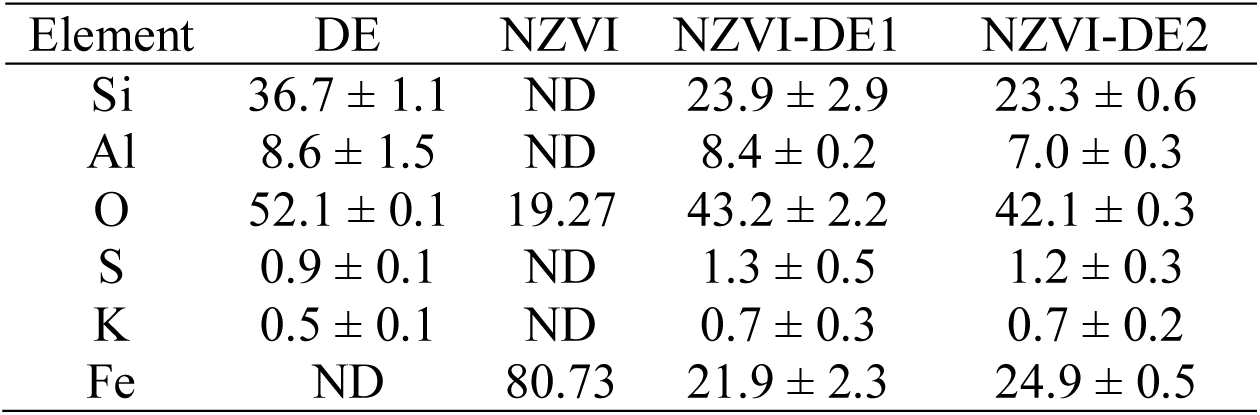
Element composition of the diverse treatments DE, NZVI, NZVI-DE1 and NZVI-DE2

| Element | DE | NZVI | NZVI-DE1 | NZVI-DE2 |
| --- | --- | --- | --- | --- |
| Si | 36.7 ± 1.1 | ND | 23.9 ± 2.9 | 23.3 ± 0.6 |
| Al | 8.6 ± 1.5 | ND | 8.4 ± 0.2 | 7.0 ± 0.3 |
| O | 52.1 ± 0.1 | 19.27 | 43.2 ± 2.2 | 42.1 ± 0.3 |
| S | 0.9 ± 0.1 | ND | 1.3 ± 0.5 | 1.2 ± 0.3 |
| K | 0.5 ± 0.1 | ND | 0.7 ± 0.3 | 0.7 ± 0.2 |
| Fe | ND | 80.73 | 21.9 ± 2.3 | 24.9 ± 0.5 |

The second most abundant element in DE is Si (37%) as was expected and elsewhere reported (Yuan et al., 2010). The Si content varies depending on the physicochemical conditions present during the formation of the DE bank (Guatame-Garcia, Buxton, 2018). These minor variations could affect the performance of the material because the higher the content of Si the more silanol groups available to react with polar organic compounds increasing the number of compounds adsorbed by DE (Al-Ghouti et al., 2003). Carbonate minerals can also affect the number of compounds that DE can absorb because the deposit of this material along the diatomite structure reduces its porosity (Guatame-Garcia, A. Buxton, 2018). However, the XRD analysis showed several crystal structures formed in the DE with the abundance of cristobalite, tridymite, and quartz, but anyone with carbonate which ensures the condition of the material to be a practical support to the nanoparticles (Fig. 1).

**Fig. 1.**
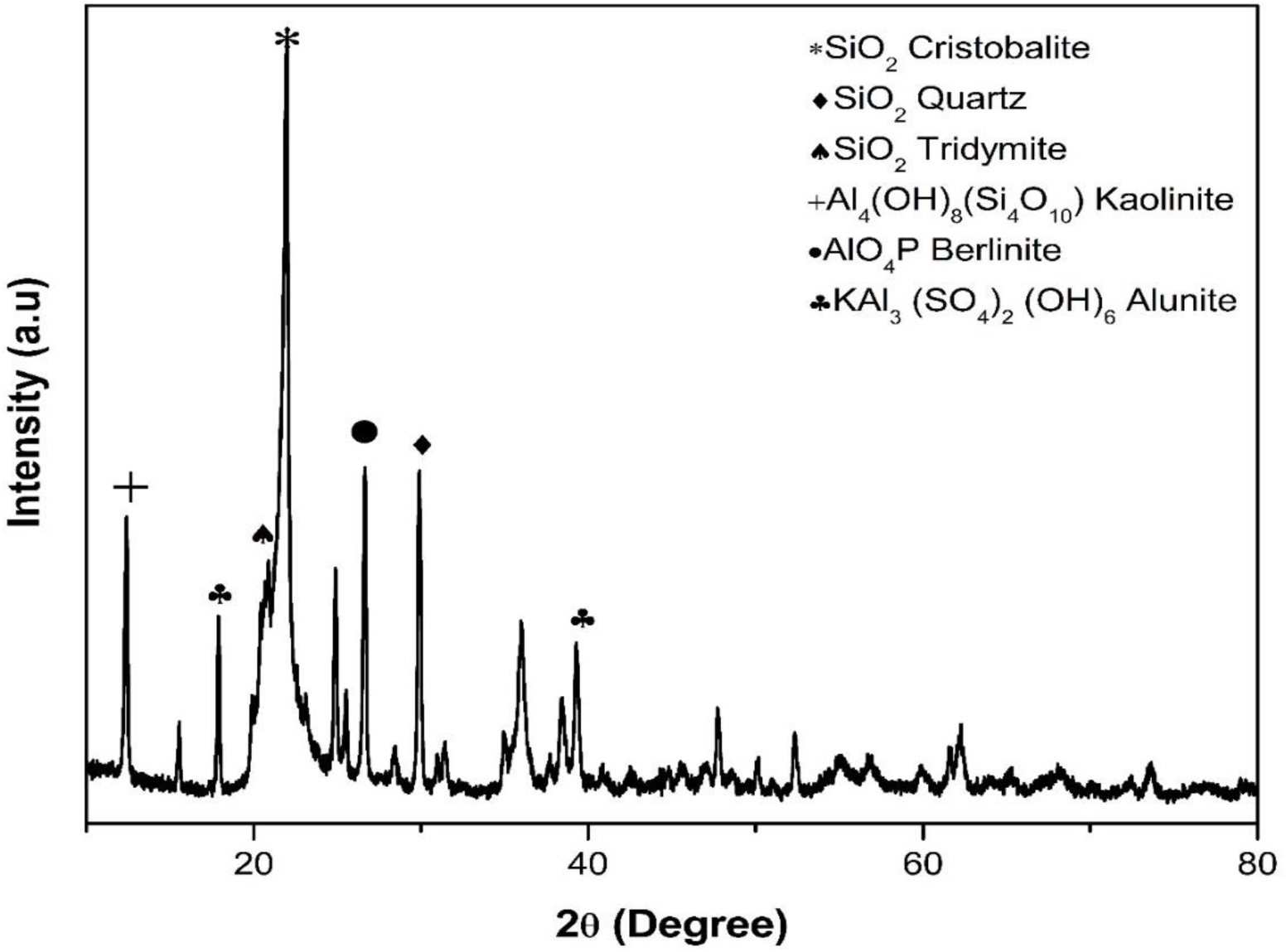
XRD pattern of diatomaceous earths

The Brunauer-Emmett-Teller (BET) specific surface area was of 23 m^2^/g with a pore volume of 0.56 cm^3^/g. As has been recorded by other DE characterization studies (Crane and Sapsford, 2018). The homogenous pore volume allows the molding of narrow size particles dispersed homogeneously on their internal surfaces (Machado et al., 2019).

Regarding NZVI-DE-1 and NZVI-DE-2, the SEM/EDX analysis showed a decrease in the levels of Si and O compared with the DE analysis given that space is occupied by iron 22 % and 25 % for NZVI-DE-1 and NZVI-DE-2, respectively (Table 1). The XRD pattern of NZVI-DE showed diffraction peaks at the 2θ of 44.90° confirming the presence of zerovalent iron in both treatments of NZVI-DE as well as on NZVI treatment (Fig. 2).

**Fig. 2.**
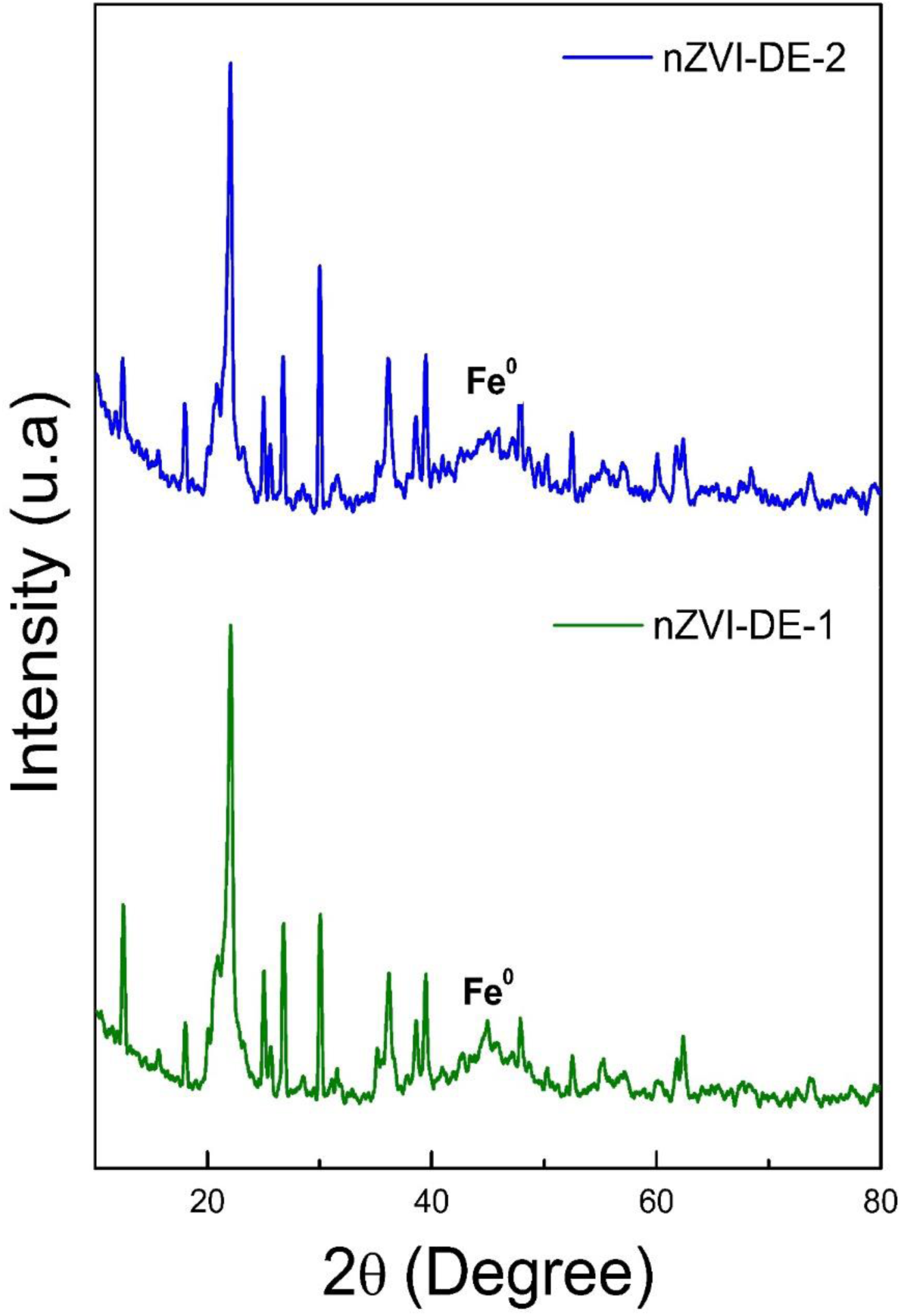
XRD pattern of nZVI-DE-1 and nZVI-DE-2 treatments

Crane and Sapsford et al. have reported similar XRD patterns of iron and DE. However, they indicate the presence of other metals such as Al, suggesting that perhaps the duration of the acid wash ∼ 2 h was not enough (Crane and Sapsford, 2018). The TEM images showed spherical forms in clusters between 20 to 40 nm, with an average size of 35 ± 8 nm (Fig. 3).

**Fig. 3.**
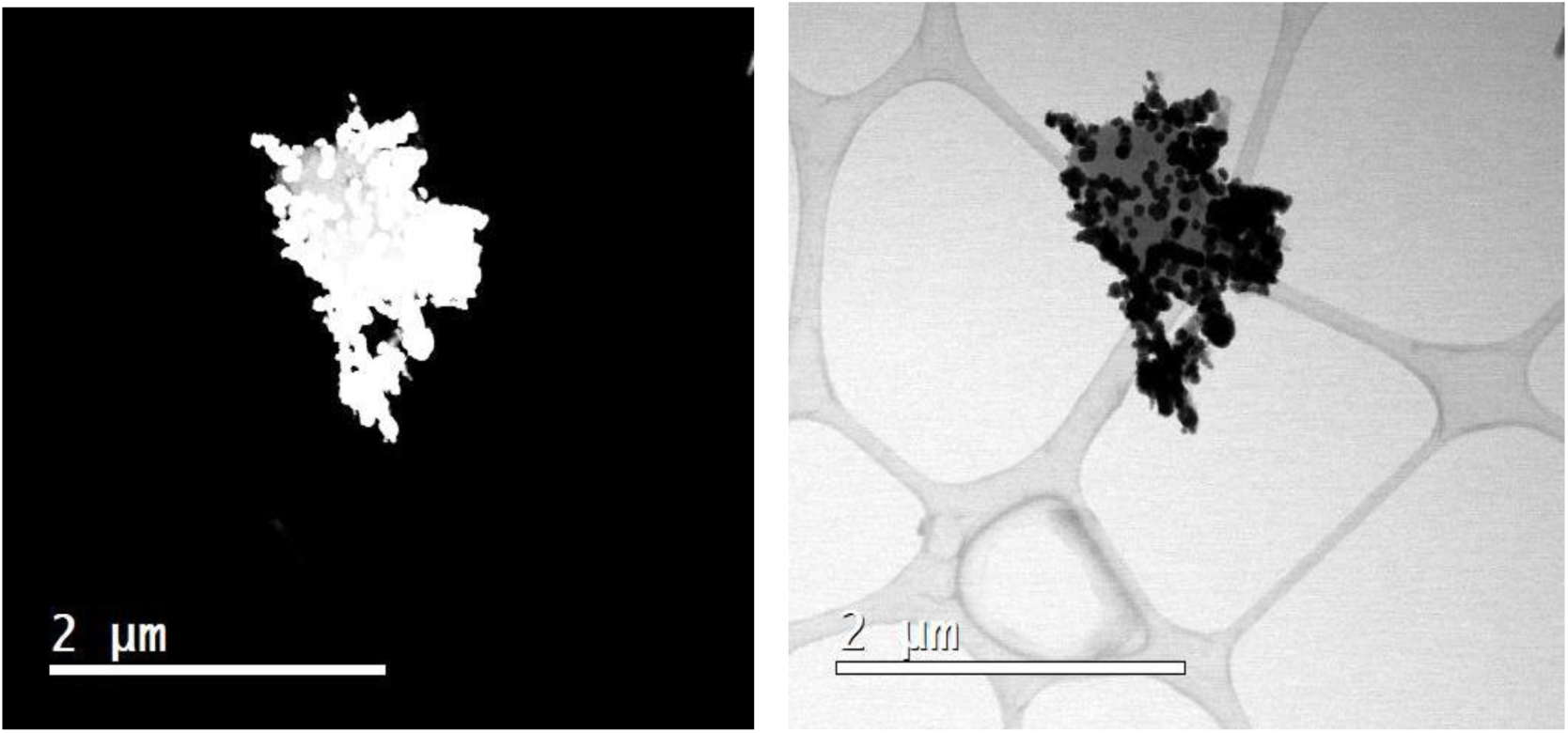
Dark field (A) and bright field (B) micrographs of the Fe nanoparticles decorating DE particles. Fe nanoparticles brighter on dark field contrast with those of the bright field

### Batch degradation experiments

The reaction proceeded in the first 5 min with a drastic change of color, and later, the difference decelerated and stopped at minute 7. Fig. 4 shows the UV-vis spectra taken 10 min. after, when the reaction was stable. The black line shows the spectra of the water with 1 g/L of AB, the red line indicates the NZVI-DE-1 treatment which removed 90% of the pigment, and the blue line shows that the NZVI-DE-2 treatment achieved 98% removal of AB. The NZVI control removed just 40 % of AB, and DE-1 and DE-2 controls removed 37% and 24% respectively. The contrast between the catalytic power of supported and unsupported nanomaterials is significant NZVI-DE-1 and NZVI-DE-2 removed 50 and 58 % more dye than unsupported nanoparticles.

**Fig. 4.**
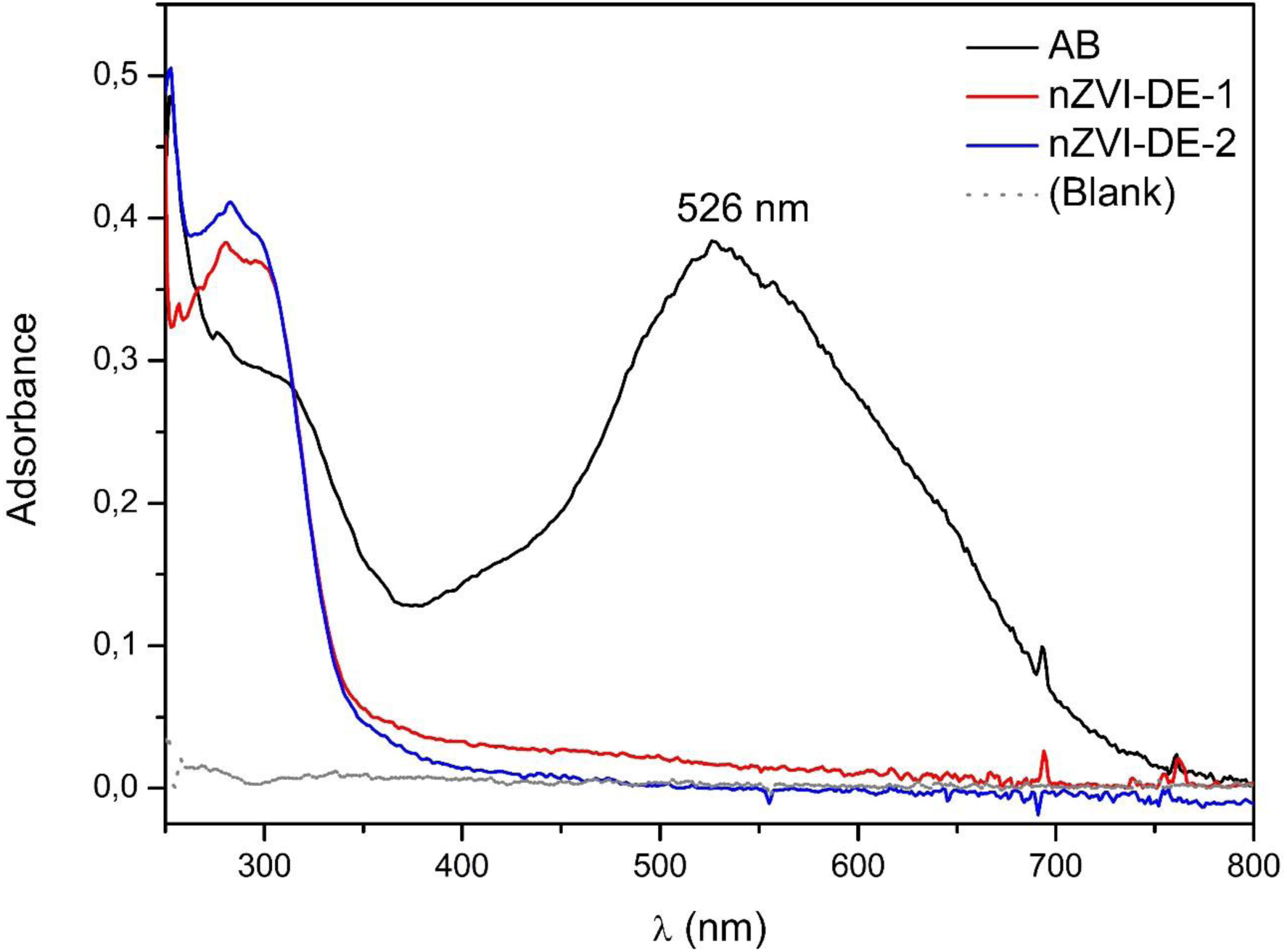
Absorbance signal of the dye and the dye removed by the NZVI-DE-1 and 2

In NZVI-DE-1 and NZVI-DE-2, the NZVI particles were responsible for the removal of the dye through reduction and adsorption processes (Deng et al., 2018; Fan et al., 2009), while the DE played a protective role for the NZVI catalyst (reducing agglomeration) and contributed to the adsorption of AB. The amount of AB removed and adsorbed could be calculated based on removal results of NZVI and DE controls, and evidently, around 40 – 50 % of the removal was adsorbed by the DE and the rest was the reduction performed by the nanoparticles. Another factor that could contrast the removal of both treatments could be the amount of DE. NZVI-DE-1 had less DE where nanoparticles were supported; and this treatment also had less iron than NZVI-DE-2. The removal performed by the nanoparticles in contact with the contaminant occurs because the nanoparticles play a role as electron mediator; also, H atoms are generated and this causes the break of the double bond (-C = C-) eliminating the chromophore group of the dye (Chen et al., 2011). Destroying one of the critical components of the dye and, as a result, the absorption peak at a wavelength of 526 nm was reduced (Fig. 4). At the same time, Fe° reacted and formed oxides such as FeO_2_, FeO3, and FeOH which have a high capacity to absorb some molecules of the pollutant (Wu et al., 2015). However, this capacity to absorb could be affected by internal changes in pH (Shah et al., 2015). The pH of the dye solution plays an essential role in the entire adsorption process and the adsorption capacity, influencing the loads of the NZVI-DE and DE. This changes the degree of ionization and dissociation of the functional groups in the active sites of the adsorbent materials (Basheer, 2018). At the beginning, the pH of the treatments was 7.4, however at the end of the removal experiments, NZVI-DE-1 had 8.9; and NZVI-DE-2 had 8.7; while NZVI had a more neutral pH of 7.7; and DE-1 and 2 had an acid pH of 4 and 4.6., respectively. The treatment with nanoparticles showed pH values around 8 because the dye removal is a dynamic process where Fe° nanoparticles begin to be transformed into oxides as Fe^2+^, Fe^3+^, Fe(OH)_3_ and Fe(OH)_2_, which reduces the quantity of H+ and raises the pH of the liquid (Garg et al., 2018). Whereas in the treatment with only DE, the pH of the solutions finishes acidic due to the protonation of surface silanol groups where protons are forming conjugate acids that lower the pH (Lowe et al., 2015; Nosrati et al., 2017).

### Toxicological analysis of treated water on zebrafish embryos

Embryos were treated with AB, and the water treated with NZVI-DE-1, NZVI-DE-2, DE-1, and DE-2 during 24, 48, 72, and 96 hours post-fertilization (hpf). Embryos exposed only to AB died after treatment; meanwhile, embryos treated with NZVI-DE-1 and NVZI-DE-2 developed alongside the embryos in control water (Fig. 5 A-C and G). Embryos grown in the presence of DE developed regularly (data not shown). The embryos treated with NZVI-DE-1 and NZVI-DE-2 were allowed to develop until 24 hpf; embryos developed normally, but a slight developmental delay was observed compared to the control (Fig.5 D-F). However, after 24 hours of culture particles associated with the chorion were visible in NZVI-DE-1 and NZVI-DE-2 samples, possibly due to an agglomeration of nanoparticles (Fig. 5 B and C asterisk), and perhaps this caused the developmental delay observed in these samples. At 48 and 72 hpf (data not shown) there was no difference in the viability of the embryos when compared to the control (Fig 5 N); but a unambiguous effect in the morphology of the embryos was observed as all of the embryos had cardiac edema, smaller eyes and curved and smaller bodies with less pigmentation (Fig. 5 H-O). We also observed particles attached to the chorion (Fig 5 B and C; A and J); these could be the nanoparticles since we do not observe these aggregates in the control embryos. At 96 hpf the embryos treated with the water from NZVI-DE-1 and NZVI-DE-2 died; meanwhile control embryos developed normally. The delay in development at 24 hpf; the morphological effect at 48 and 72 hpf and the comprised viability of embryos at 96 hpf could be due to the presence of the nanoparticles. Previous work done by Zhu X et al. (2012) showed that embryos exposed to iron nanoparticles caused effects in survival and morphological defects similar to what we observe, such as cardiac edema.

**Fig. 5.**
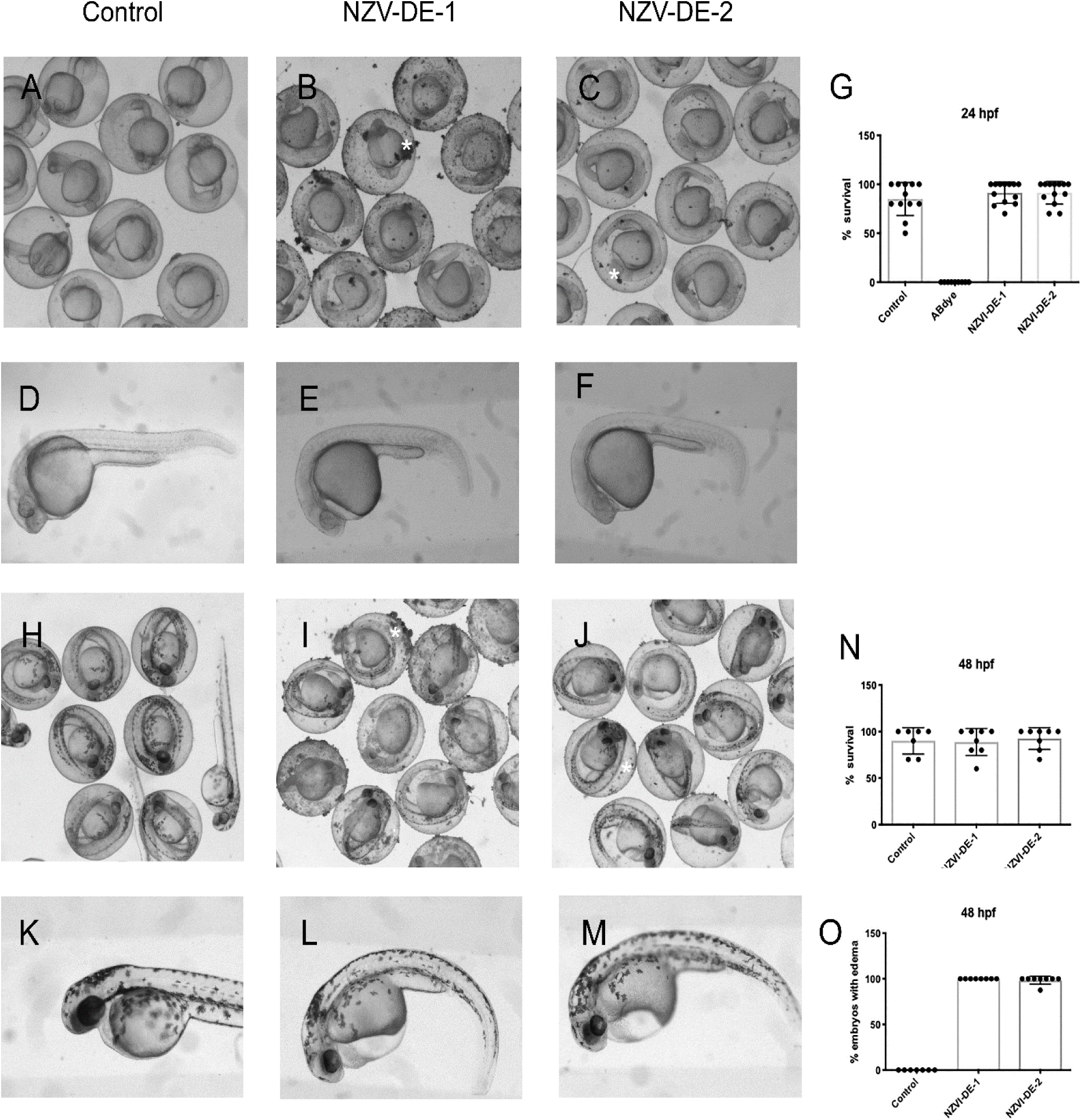
Zebrafish embryonic development progress in the presence of NZV-DE-1 and NZV-DE2. **A**. Control embryos at 24hpf. **B and C**. Embryos at 24hpf proceed with development when exposed to contaminated water is treated with NZV-DE. Agglomeration of nanoparticles can be observed in the chorion (asterisk in B and C). **D-F**. A slight delay in development is observed in NZV-DE-1 (E) and NZV-DE-2 (F) embryos compared to the control (D). **G and N** Viability is fully restored in NZV-DE treated water exposure compared to AB dye exposure. **H** Control embryos at 48 hpf. **I and J** Embryos developed until 48 hpf as the control embryos; nevertheless, the embryos treated with NZV-DE (L and M) present cardiac edema (O), smaller eyes curved body plan, less pigmentation and are smaller compared to the control (K).

Nevertheless, the improvement of embryonic viability was highly significant compared to the embryos treated only with the AB dye (Fig.6). In order to determine the exact moment that the AB had an effect, dye cultures of 4 hours were carried out. Here we could observe that after 4 hours of treatment all of the embryos treated with AB dye were already dead; meanwhile control embryos and those treated with NZV1-DE-1 and NZV1-DE-2 developed normally (Fig 6). Additionally, the embryos treated with AB dye were colored as was the chorion (Fig. 6D and H), while NZV1-DE-1 and NZV1-DE-2 treated embryos followed normal development without coloration (Fig. 6). Our results suggest that the treatment of NZV1-DE-1 or NZV1-DE-2 is sufficient to avoid the lethality observed in embryos exposed to AB dye given that development proceed normally at the first hours of treatment. Nevertheless, further stages of development, such as 48 and 72 hpf, were affected as a result of the nanoparticles present in the samples. The effects observed in the embryos after 24 hours could be mitigated by potential applications of NZVI-DE via filters where the treated water is in contact with the organism for a short period of time

**Fig. 6.**
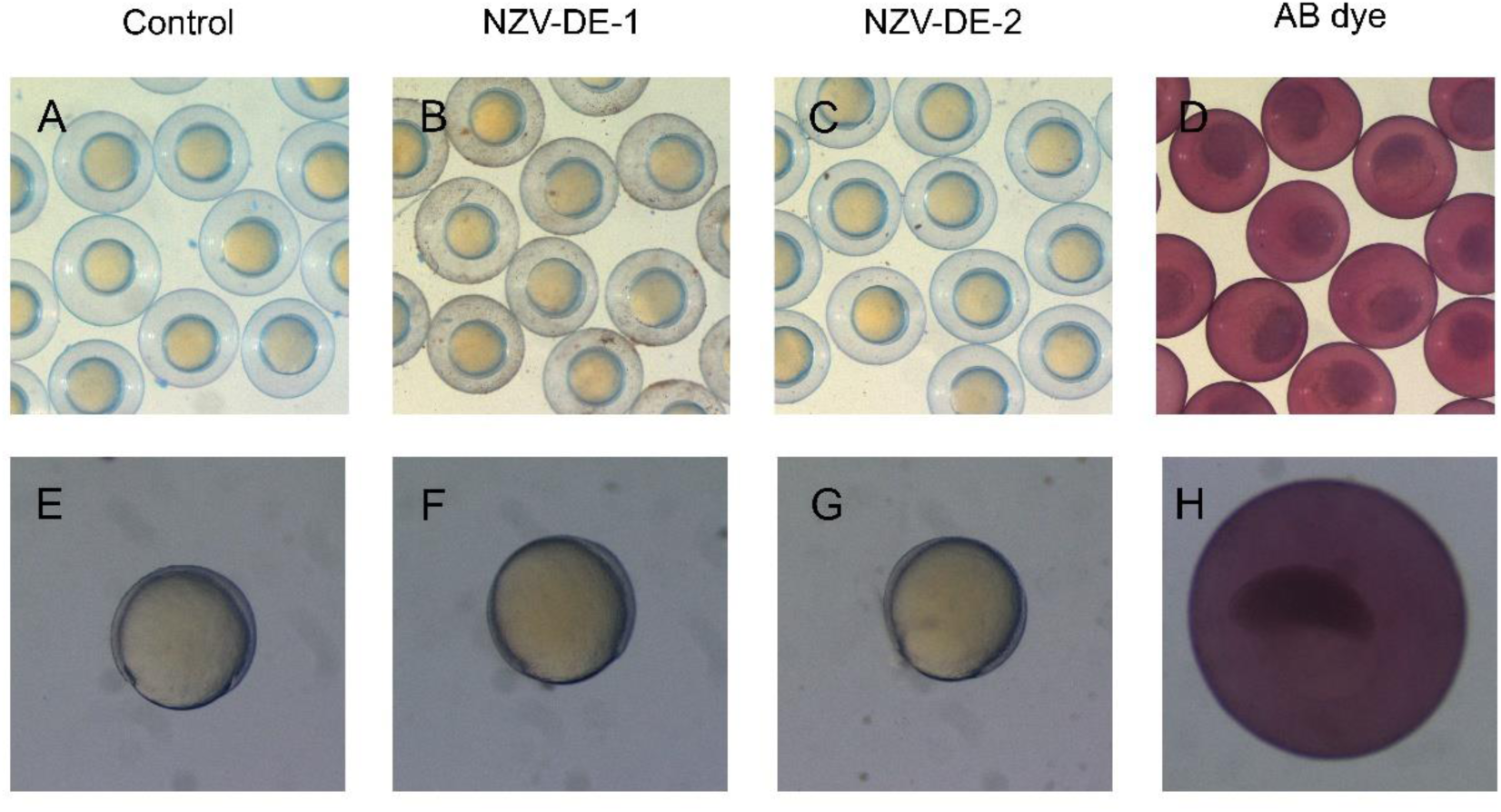
Early zebrafish development is restored after NZV-DE treatment. **A**. Control embryos after 4 hours of treatment. **B** Embryos treated with NZV-DE-1 develop normally. **C**. Embryos treated with NZV-DE-2 develop as the control embryos. **D** Embryos treated with AB dye died after 4 hours of treatment. **E-H** Higher magnifications show that embryos treated with NZV-DE (F and G) develop as the control embryos (E); meanwhile embryos treated with AB dye absorb the dye and do not develop further and die.

## Conclusion

There is increasing interest in the use of nanoparticles to remove pollutants, however the possible risk to aquatic life or through food chains are not sufficiently clear. In this experiment, the NZVI-DE-1 and NZVI-DE-2 successfully removed more than 90 % of one of the most widely used pollutants, acid blue. The toxicology test of the treated water showed that wild-type zebrafish (*Danio rerio*) embryos developed completely normally during the first 24 h. However, after 48 h. all of the embryos had cardiac edema, smaller eyes, and smaller bodies with less pigmentation compared to the control sample. Consequently, we suggest that nanoparticles technologies use supported materials and that nanoparticle technologies used to remove pollutants have limited contact with streams in order to avoid adverse effects for aquatic life.

## Acknowledgements

The authors thank ZEOLITECH for the donation of diatomaceous earth, Dr. Daniel Bahena Uribe and Dr. Jorge Roque De La Puente of LANE, CINVESTAV, for their excellent technical help and advice on the characterization of nanoparticles. Also, LB-D thanks the Consejo Nacional de Ciencia y Tecnología (CONACYT) and their program CATEDRAS for their support of the Project 285.

